# PxBLAT: An efficient python binding library for BLAT

**DOI:** 10.1101/2023.08.02.551686

**Authors:** Yangyang Li, Rendong Yang

## Abstract

We introduce PxBLAT, a Python library designed to enhance usability and efficiency in interacting with the BLAST-like alignment tool (BLAT). PxBLAT provides an intuitive Application Programming Interface (API) design, allowing the incorporation of its functionality directly into Python-based bioinformatics workflows. Moreover, PxBLAT’s design philosophy emphasizes ease of use, memory efficiency, and the elimination of intermediary files and unnecessary system calls, thereby enhancing computational speed and user experience. Benchmark tests reveal its superior performance across various datasets, illustrating its capacity to maintain correctness. PxBLAT supports Python (version 3.9+), and pre-compiled packages are released via PyPI (https://pypi.org/project/pxblat/) and Bioconda (https://anaconda.org/bioconda/pxblat). The source code and executable are freely available for academic, nonprofit, and personal use. Its documentation is available on ReadTheDocs (https://pxblat.readthedocs.io/en/latest/).

## Introduction

The rise of Python as a preferred programming language within bioinformatics is widely acknowledged as a result of its user-friendly nature, extensive libraries, and unparalleled versatility [1]. A variety of libraries have been crafted to augment Python’s interface, thereby amplifying the adaptability and compatibility of bioinformatics tools [2, 3]. For instance, Biopython [2], a preeminent bioinformatics library, furnishes interfaces to tools like Basic Local Alignment Search Tool (BLAST) [4] and Clustal [5].

Furthermore, the unprecedented growth in genome sequencing technologies has significantly increased the availability of genomic data, emphasizing the need for advanced tools in both research and clinical contexts. BLAT [6], a prominent tool in bioinformatics, is renowned for its speed in genome sequence alignments and serves as a more efficient alternative to BLAST [4] for aligning DNA sequences with the human genome. However, despite its popularity and effectiveness, BLAT’s integration is fraught with difficulties, primarily due to its C-based implementation and reliance on Command-Line Interfaces (CLIs), hindering seamless integration into Python projects. Moreover, BLAT’s reliance on system calls and data conversions introduces considerable overhead. Until recently, a comprehensive solution to effectively enhance BLAT’s usability within Python frameworks was missing.

Also, executing extensive queries with the BLAT suite often leads to inefficiencies, particularly when operations are isolated and not executed in batches. Typically, BLAT’s task allocation is sporadic, intermixed with other unrelated tasks. Users generally face a choice: either employ standalone BLAT or integrate *gfServer* with *gfClient*. BLAT’s standard operational model involves initiating *gfServer*, conducting the sequence query through *gfClient*, and subsequently terminating the server after each query. This method becomes highly inefficient for ungrouped, numerous queries as it necessitates the repeated initialization and shutdown of *gfServer*, introducing significant overhead. This overhead is particularly evident in tasks such as indexing large references, where, for instance, indexing the human genome (hg38) alone consumes approximately five minutes. An optimized approach would entail initiating *gfServer* a single time and leveraging *gfClient* to execute multiple queries. However, the command-line-only access to *gfServer* and *gfClient* complicates this process. This limitation necessitates the management of system calls (like *subprocess* or *os*.*system*), the handling of intermediate temporary files, and dealing with format conversion, all of which cumulatively degrade performance.

PxBLAT is proposed as a solution that allows for the programmatic use of BLAT, ensuring its smooth integration into new algorithms or analytical pipelines within the Python ecosystem. It acts as a conduit, merging the high-performance capabilities of BLAT with Python’s versatility while ensuring data reproducibility. The primary goal of PxBLAT is to bridge the gap in the current landscape by providing a Python binding library tailored specifically for BLAT, addressing both the efficiency bottlenecks and the ergonomic challenges of its integration.

## Design and implementation

The design of PxBLAT is anchored in the principles of readability and simplicity, fostering an intuitive user interface that minimizes the learning curve for users. In our quest to streamline complexity and amplify both usability and performance, we meticulously extracted the core implementation of BLAT from the broader UCSC Genome Browser (UCSC) codebase, significantly reducing dependency overhead.

We preserved the integrity of the original C codebase while reimplementing key BLAT (V37.1) utilities such as *faTwoBit, gfServer*, and *gfClient* in C`++`. This strategic choice not only modernizes the code but also enhances maintainability and scalability. The integration of the revamped C`++` code with PxBLAT was achieved using Pybind11 [7], a lightweight, seamless method for interfacing C`++` and Python.

This approach ensures a direct and efficient interaction with BLAT’s functions, upholding the original performance benchmarks and reliability of BLAT. Simultaneously, it extends the framework’s functionality, aligning it with modern computational standards and making it a robust tool in the bioinformatics toolkit (Table 1).

**Table 1.** Overview of features of PxBLAT compared with BLAT. This table presents a comprehensive comparison between the features offered by PxBLAT and BLAT. Features are denoted with a ✓ to signify availability and an ✗ to indicate absence. Notably, PxBLAT showcases significant enhancements, particularly in server management (e.g., wait server), data conversion (e.g., fasta to bit, bit to fasta), and enriched user interaction (e.g., shell completion). These advancements firmly establish PxBLAT as a superior and more versatile alternative to the conventional BLAT tool.

| Feature | PxBLAT | BLAT |
| --- | --- | --- |
| start server | ✓ | ✓ |
| stop server | ✓ | ✓ |
| query server | ✓ | ✓ |
| wait server | ✓ | ✗ |
| fasta to bit | ✓ | ✗ |
| bit to fasta | ✓ | ✗ |
| port retry | ✓ | ✗ |
| shell completion | ✓ | ✗ |

PxBLAT delivers its query results in alignment with the *QueryResult* class of Biopython [2], enabling seamless manipulation of query outputs using Biopython’s comprehensive suite of features (Listing 1). This integration effectively streamlines the post-query workflow, allowing users to leverage the full potential of Biopython in their sequence alignment tasks. Significantly, PxBLAT negates the necessity for intermediate files by conducting all operations in memory. This advancement eliminates the often cumbersome and time-consuming step of data format conversion, enabling users to concentrate on the core aspects of sequence alignment. To enhance user flexibility, the necessity for input and output files has been made optional, aligning with diverse user preferences and workflows.

Recognizing the latency and potential performance bottlenecks induced by system calls, PxBLAT minimizes their usage, thereby streamlining operations and enhancing overall efficienc. Additionally, PxBLAT simplifies server status retrieval, circumventing the complexities and potential pitfalls of log file manipulation, particularly in concurrent usage scenarios. To further elevate the user experience and operational efficiency, PxBLAT integrates several ergonomic features. These include real-time server readiness checks for alignments, automatic port retries when the default is in use, and the capability to latch onto an already running server if available. These features collectively ensure a smoother, more efficient alignment process, reducing downtime and maximizing productivity.

To facilitate a smooth user onboarding experience, we offer an extensive range of examples and comprehensive documentation (Listing 1). PxBLAT introduces a robust set of APIs, including the classes *Server* and *Client*, along with a suite of functions designed to replicate the capabilities of the BLAT suite. These classes mirror the utilities of the CLI tools *gfServer* and *gfClient*, respectively, but with added flexibility to accommodate a wider range of user requirements. Key functions such as *start server, query server, status server, fa2twobit*, and *twobit2fa* are provided to cater to diverse usage scenarios. Rigorous testing and development protocols, incorporating Continuous Integration (CI) and Continuous Development (CD), have been employed to ensure high code quality and reliability.

Moreover, PxBLAT utilizes type annotations in its public classes and functions. This not only reinforces code quality and correctness through type checking and static analysis but also enhances the development experience. The annotated types facilitate automatic suggestion and correction of function signatures in development environments, streamlining the coding process.

In addition to the APIs, PxBLAT features CLI utilities crafted through its APIs, boasting shell completion for various systems to augment its versatility (Table 1). Recognizing the diverse technological landscape, we provide the library in wheel format compatible with multiple platforms, including Linux x86-64, macOS x86-64, and macOS arm64. This ensures a seamless installation process, free from the complexities of C library dependencies, making it straightforward and user-friendly.

**Listing 1.**
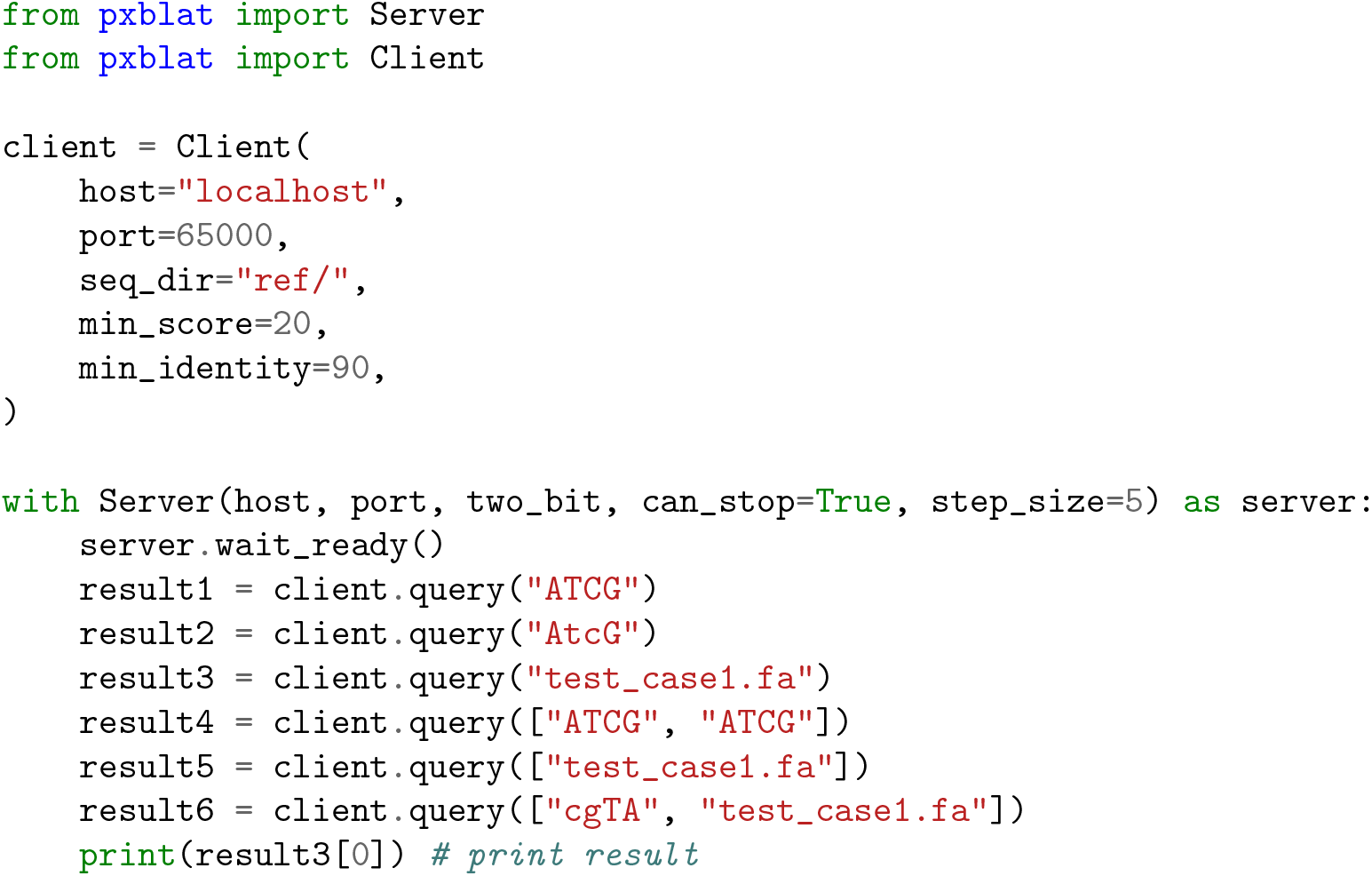
API Example. The code snippet shows how to use the API of PxBLAT, and the query result can be iterated. More code examples can be found at https://pxblat.readthedocs.io/en

## Results

### Performance on real datasets

The performance of PxBLAT were rigorously benchmarked against BLAT (V37.1) utilizing eight distinct sample sets of FASTA files. Each set comprised a group of samples, ranging from 50 to 600 samples per set. The datasets are sampled from chromosome 20 of the Human Genome (hg38), with each sample containing a single sequence. These sequences varied in length from 1000.00 bp to 3000.00 bp, encompassing a spectrum of typical use-case scenarios (Fig 1).

**Fig 1.**
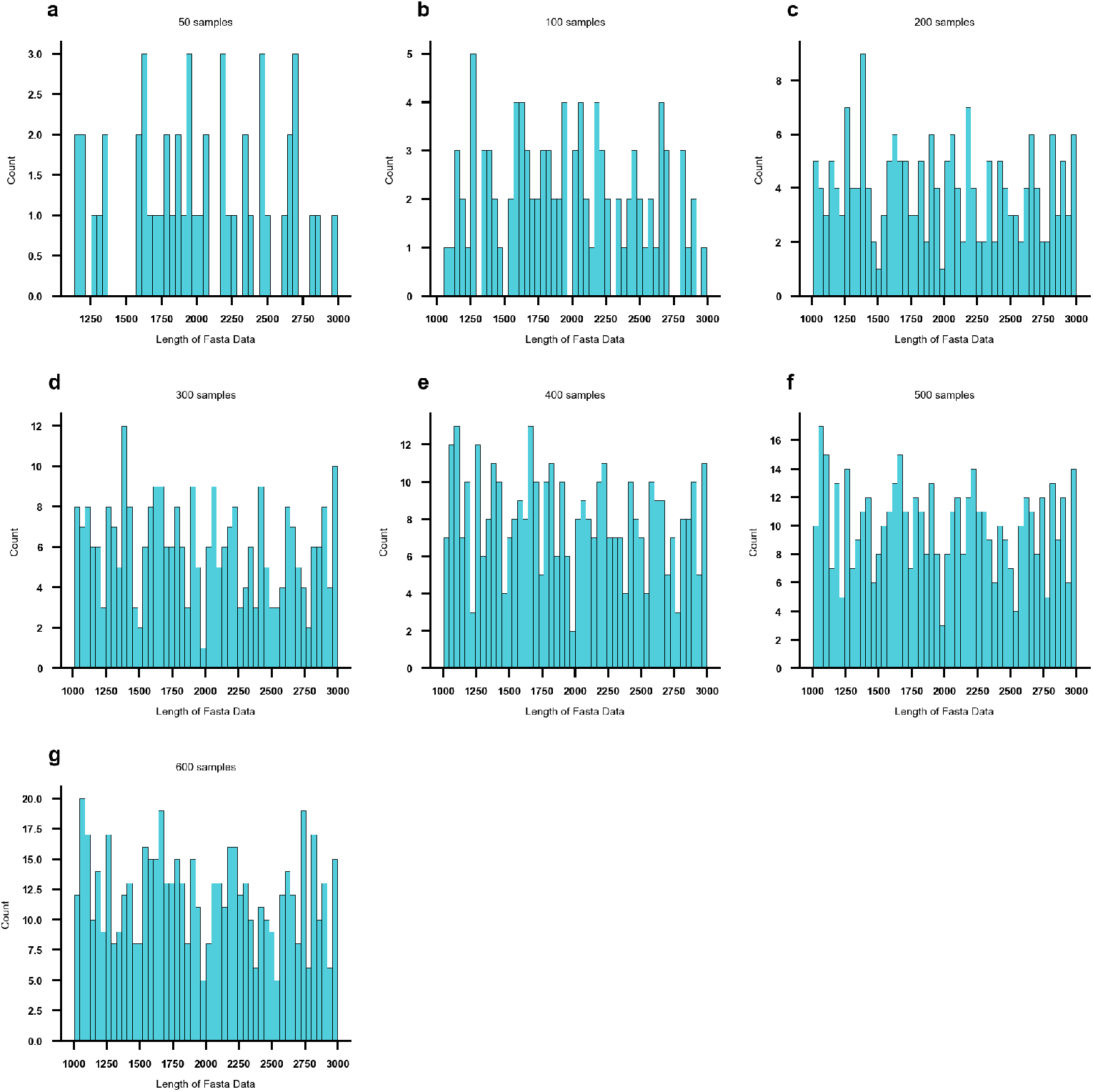
Sequence length distribution in real datasets. This figure illustrates the distribution of fasta sequence lengths across different sample sets. The *x* axis represents the sequence length, while the *y* axis denotes the count of each length. a: Distribution of a set of 50.00 samples. b: Distribution of a set of 100.00 samples. c: Distribution of a set of 200.00 samples. d: Distribution of a set of 300.00 samples. e: Distribution of a set of 400.00 samples. f: Distribution of a set of 500.00 samples. g: Distribution of a set of 600.00 samples.

To ascertain the accuracy and reliability of PxBLAT, we conducted a comparative analysis of the High-Scoring Pairs (HSPs) generated by both BLAT and PxBLAT for each sample. This comparison revealed a high degree of concordance between the HSPs produced by PxBLAT and those generated by BLAT, affirming the fidelity of PxBLAT’s output (S2 Table).

The benchmarking process was carried out on an Apple M1 Pro running macOS 13.4.1 (arm64). For launching BLAT, system calls were utilized, and the execution time was measured using the time library. Each set of FASTA files underwent three experimental runs, facilitating a comprehensive assessment of performance. The results highlighted efficiency of PxBLAT, with observed speedups ranging from 1.00 to 1.77 times compared to the BLAT execution (Fig 2).

**Fig 2.**
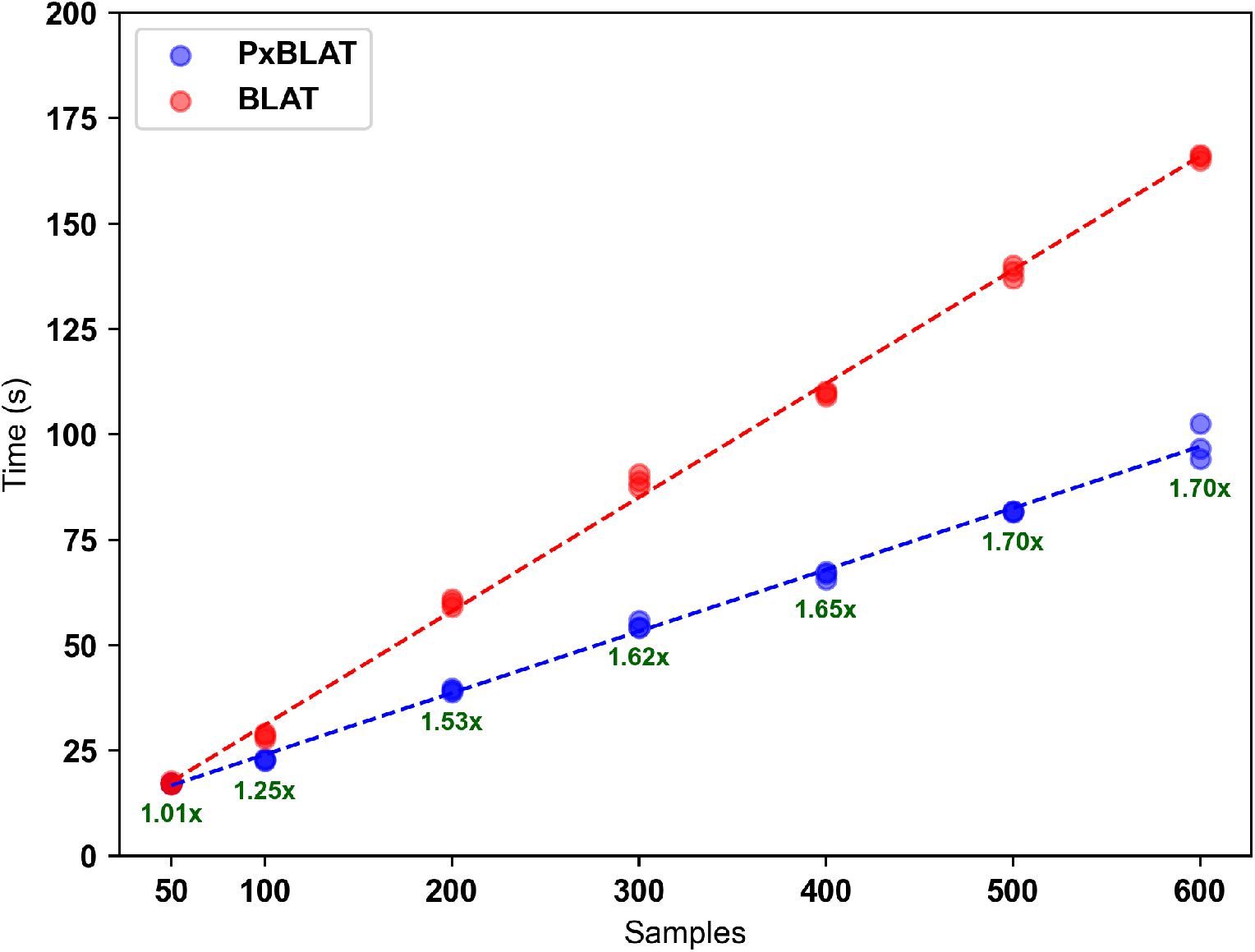
Performance comparison between BLAT and PxBLAT. This figure quantifies the performance of BLAT (indicated by red points) and PxBLAT (indicated by blue points) across various data sets, with the *x* axis categorizing the number of samples in the sets and the *y* axis detailing the execution time in seconds. Each group encapsulates the results of three independent experiments. Trend lines, depicted in red for BLAT and blue for PxBLAT, illustrate the general performance pattern for each tool. Notably, the green text highlights the speedup achieved by PxBLAT, calculated as the ratio of the execution time (time_blat_ */*time_pxblat_), underscoring the efficiency gains of PxBLAT relative to BLAT.

In summary, PxBLAT demonstrates significant advantages in terms of execution time reduction and enhanced user experience. These findings underscore its utility as a substantial improvement over the BLAT, reinforcing its value within the bioinformatics toolkit.

### Availability and future directions

The PxBLAT, along with the source code, is publicly available in the GitHub repository at https://github.com/ylab-hi/pxblat. The documentation is available at ReadtheDocs https://pxblat.readthedocs.io/en/latest/. The script for benchmarking is available at *tests/test result*.*py* in the repository. The testing dataset is available at the GitHub repository https://github.com/ylab-hi/pxblat. The path of the testing dataset is *benchmark/fas*.

As we advance PxBLAT, our vision is firmly set on developing a robust, user-oriented tool that dynamically meets the evolving demands of the bioinformatics community. Our commitment to continuous improvement is guided by insightful user feedback and the latest scientific advancements. Future enhancements of PxBLAT are directed towards augmenting functionality through the introduction of innovative features and the refinement of existing ones, thereby expanding the tool’s adaptability to a diverse range of data types and bioinformatics workflows. Simultaneously, we recognize the critical role of community engagement in shaping a tool that truly reflects user needs. To this end, we are dedicated to fostering deeper connections within the community via forums, workshops, and collaborative projects.

A relentless pursuit of performance optimization remains at the core of our efforts, aiming to significantly enhance efficiency, reduce computational overhead, and streamline the overall user experience. Additionally, we are committed to the development of comprehensive educational materials, including detailed tutorials, insightful case studies, and best practice guidelines. These resources are meticulously crafted to empower users, enabling them to fully leverage the capabilities of PxBLAT and advance their bioinformatics projects. Together, these focused endeavors epitomize our dedication to excellence and our aspiration to nurture an inclusive, innovative, and community-driven ecosystem in the field of bioinformatics.

## Supporting information

**S1 Table.** Performance comparison between PxBLAT and BLAT. This table illustrates the performance of both PxBLAT and BLAT across data sets containing 50.00, 100.00, 200.00, 300.00, 400.00, 500.00, and 600.00 samples. For each data set, three independent experiments are conducted to ensure robust performance evaluation. The efficiency of PxBLAT relative to BLAT is quantified through the speedup metric, calculated as the ratio of the execution time (time_blat_ */*time_pxblat_). This comparison highlights the computational advantages of PxBLAT in terms of processing speed.

| Samples | PxBLAT (s) | BLAT (s) | Speedup |
| --- | --- | --- | --- |
| 50.00 | 17.10 | 17.03 | 1.00 |
| 50.00 | 17.09 | 17.01 | 1.00 |
| 50.00 | 17.10 | 17.72 | 1.04 |
| 100.00 | 22.75 | 28.61 | 1.26 |
| 100.00 | 23.12 | 29.10 | 1.26 |
| 100.00 | 22.53 | 27.90 | 1.24 |
| 200.00 | 39.31 | 59.03 | 1.50 |
| 200.00 | 38.86 | 60.20 | 1.55 |
| 200.00 | 39.80 | 60.87 | 1.53 |
| 300.00 | 55.89 | 90.68 | 1.62 |
| 300.00 | 54.34 | 88.88 | 1.64 |
| 300.00 | 54.15 | 87.46 | 1.62 |
| 400.00 | 65.50 | 110.20 | 1.68 |
| 400.00 | 67.02 | 109.06 | 1.63 |
| 400.00 | 67.46 | 109.79 | 1.63 |
| 500.00 | 81.56 | 140.13 | 1.72 |
| 500.00 | 81.77 | 138.66 | 1.70 |
| 500.00 | 81.73 | 137.02 | 1.68 |
| 600.00 | 96.62 | 165.87 | 1.72 |
| 600.00 | 102.55 | 164.93 | 1.61 |
| 600.00 | 94.21 | 166.38 | 1.77 |

**S2 Table.**
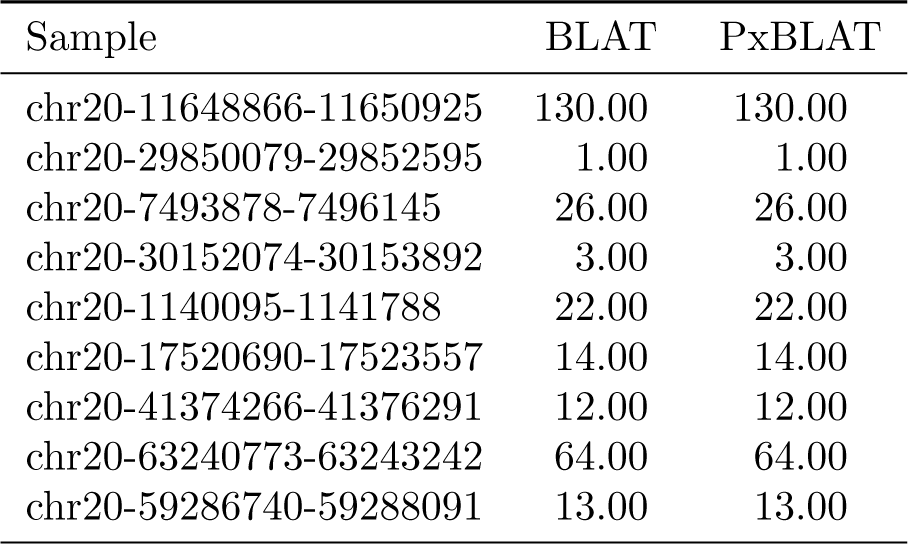
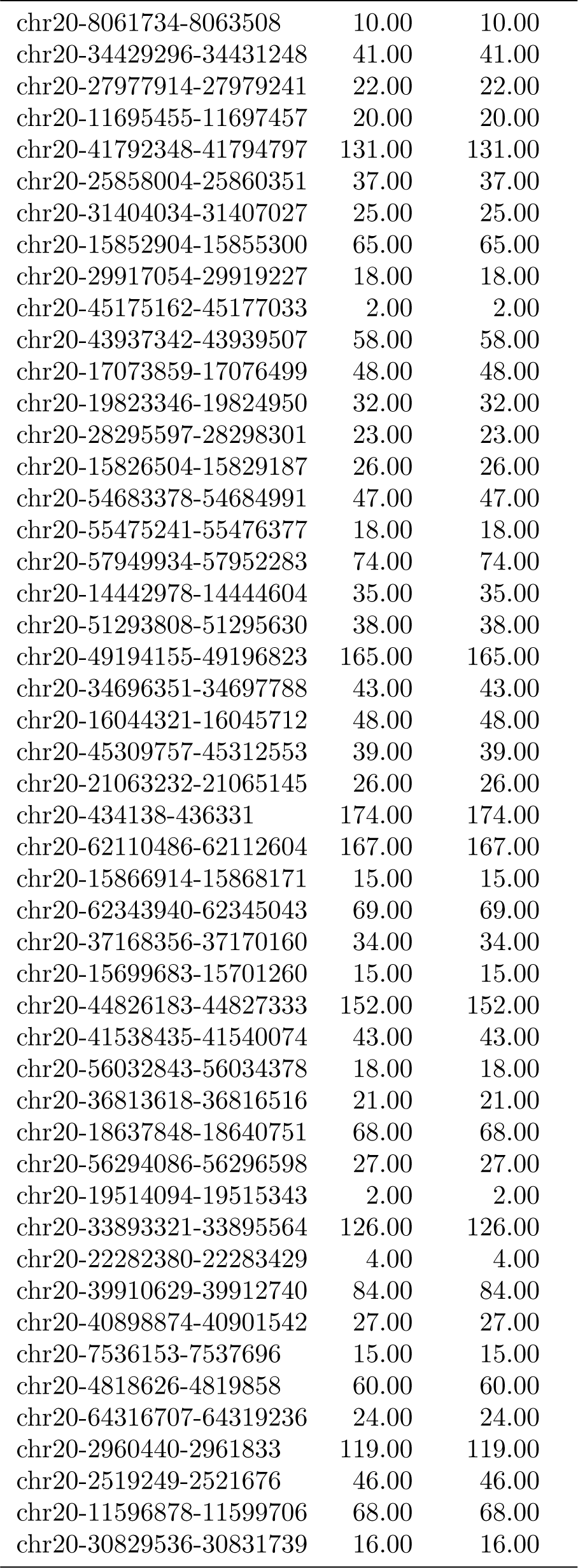
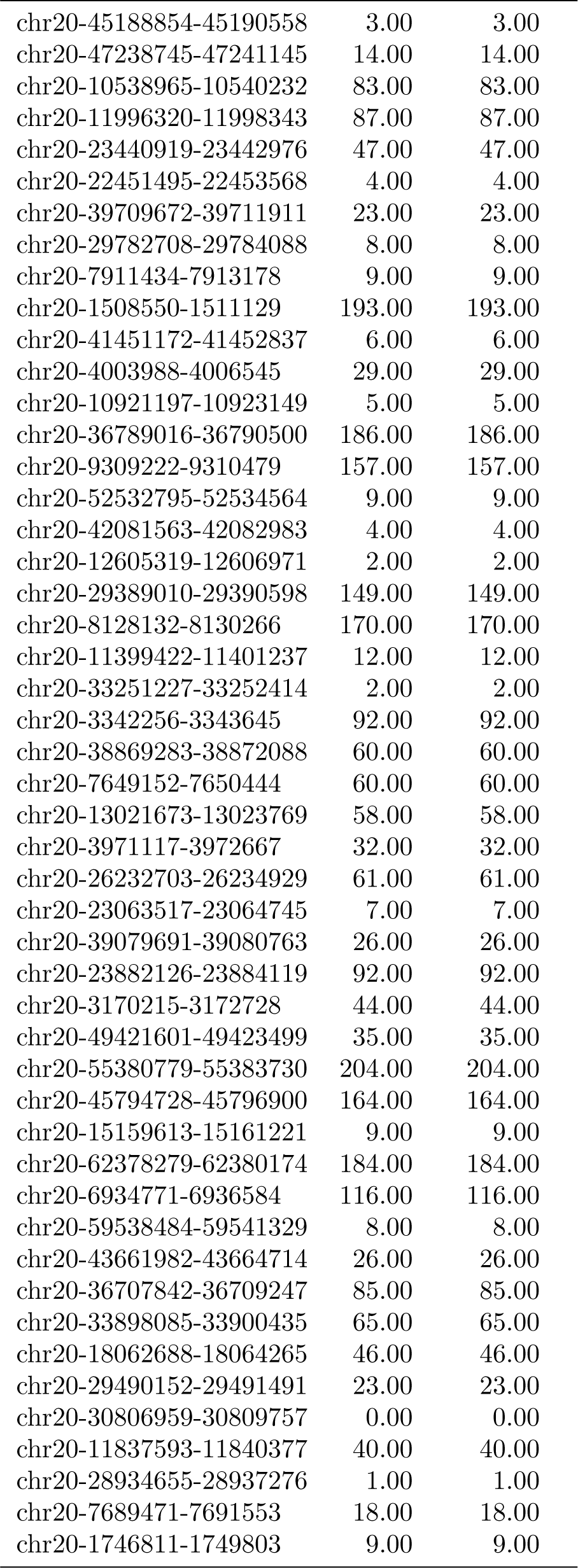
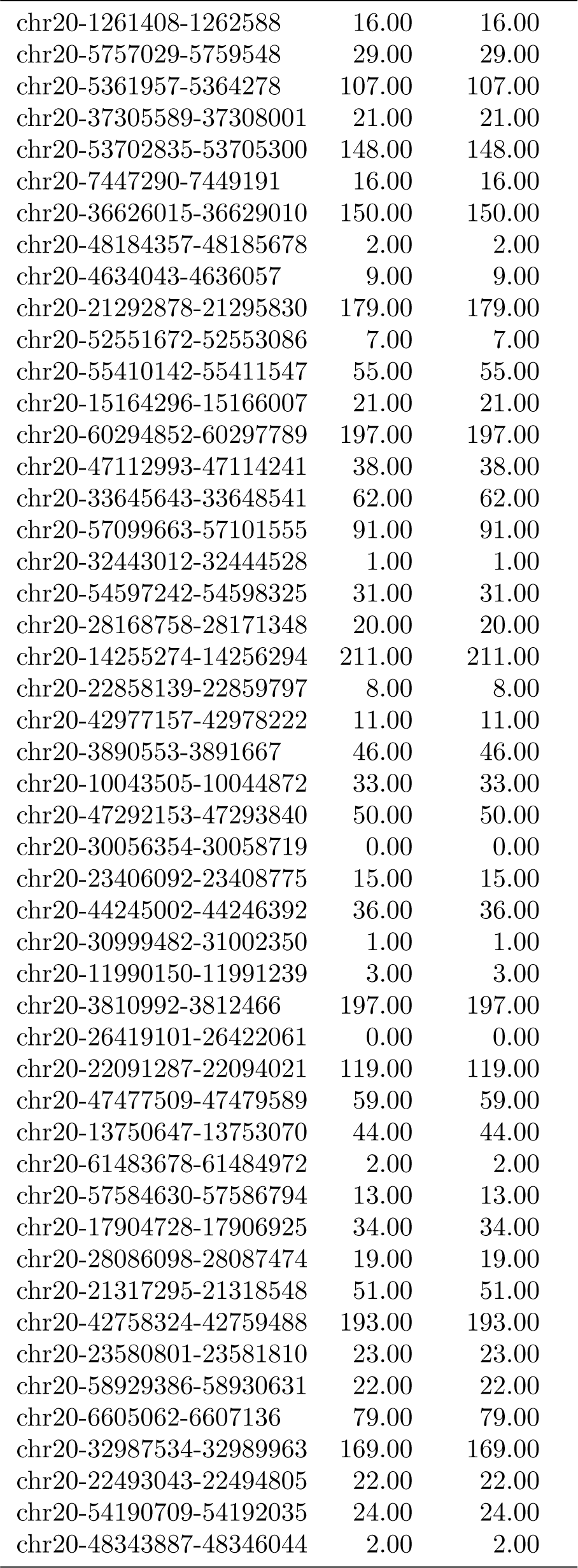
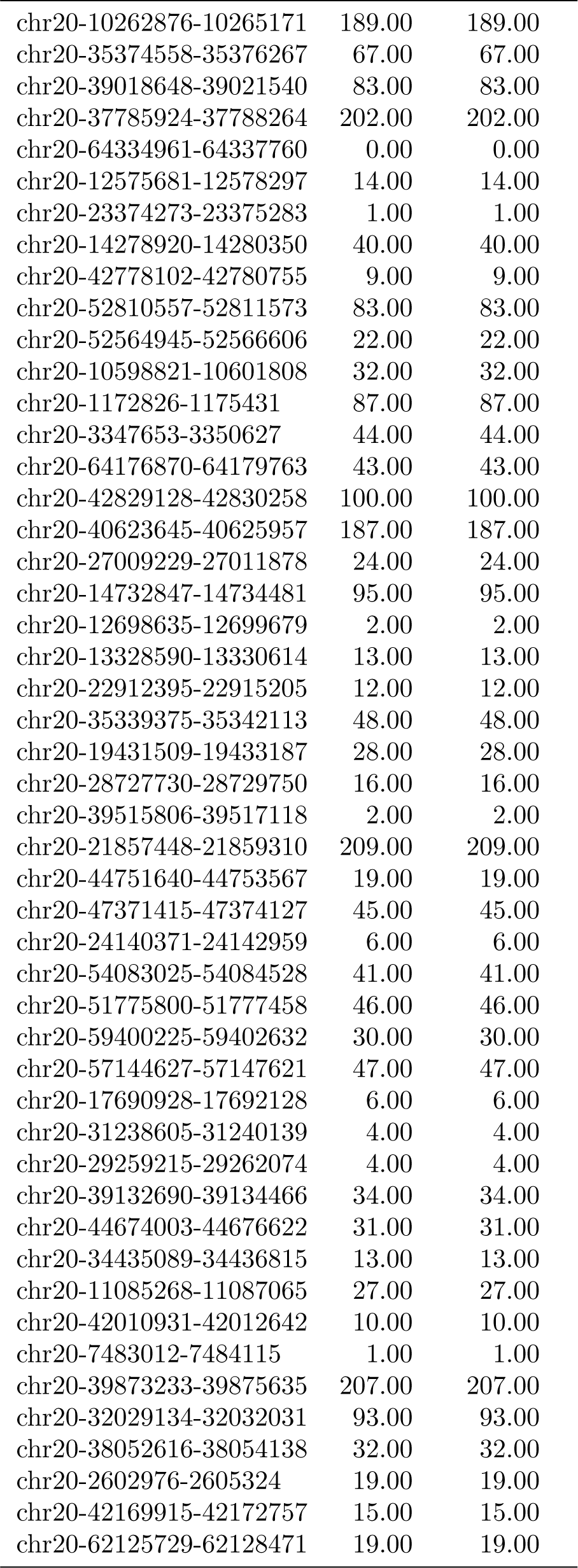
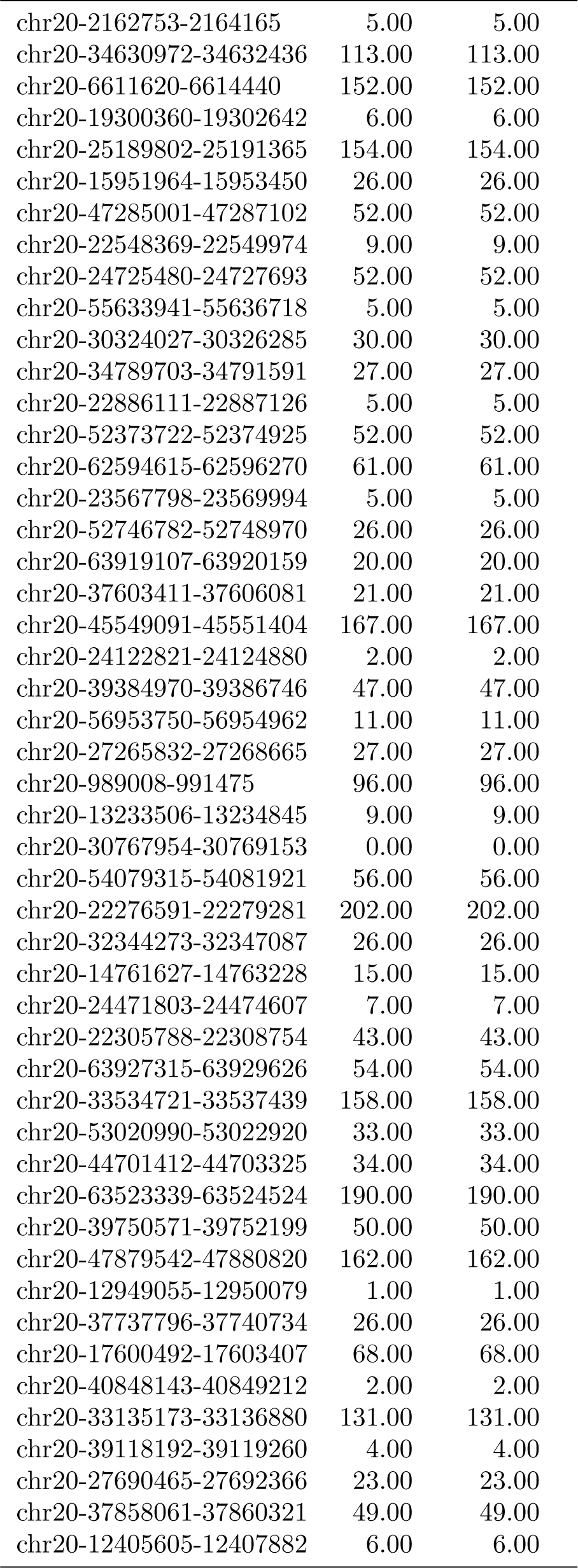
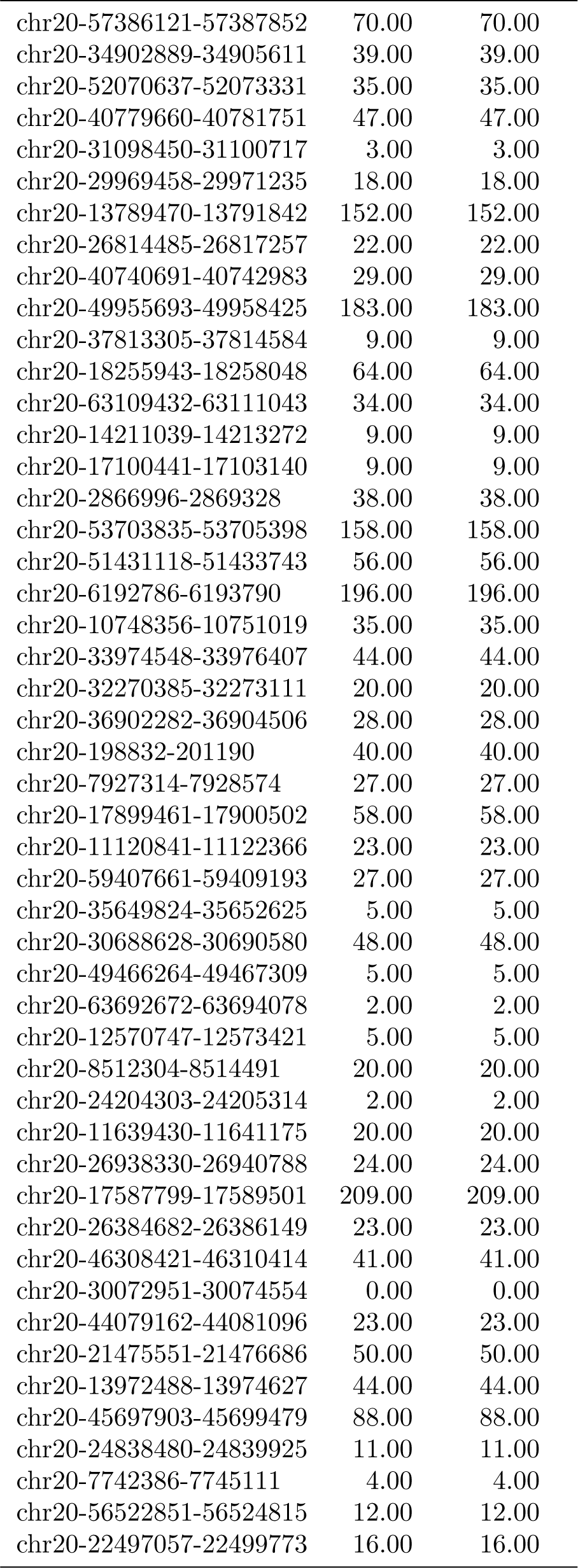
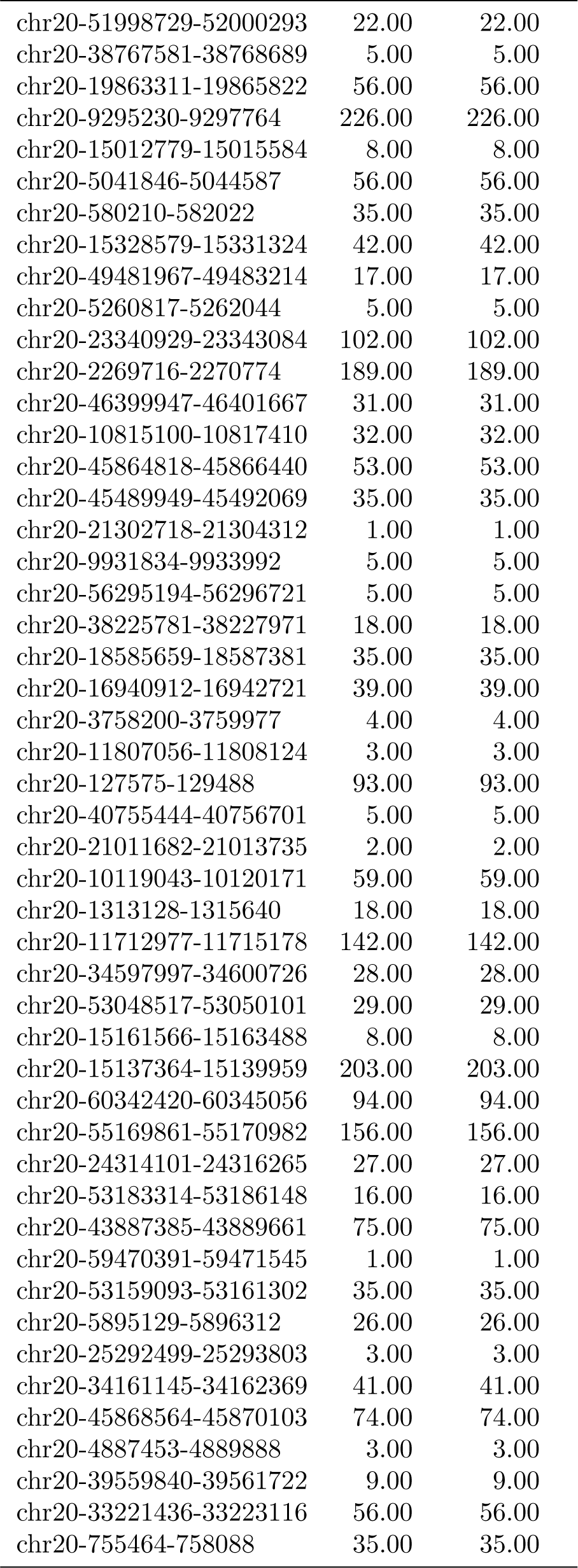
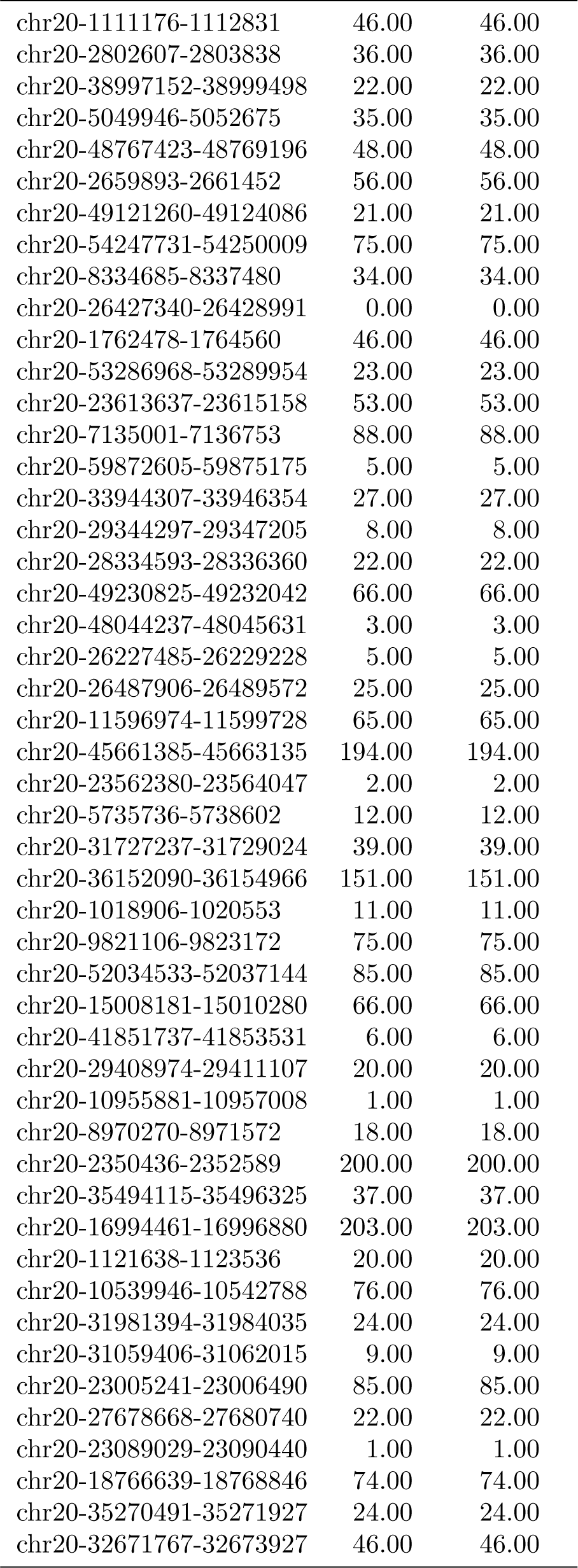
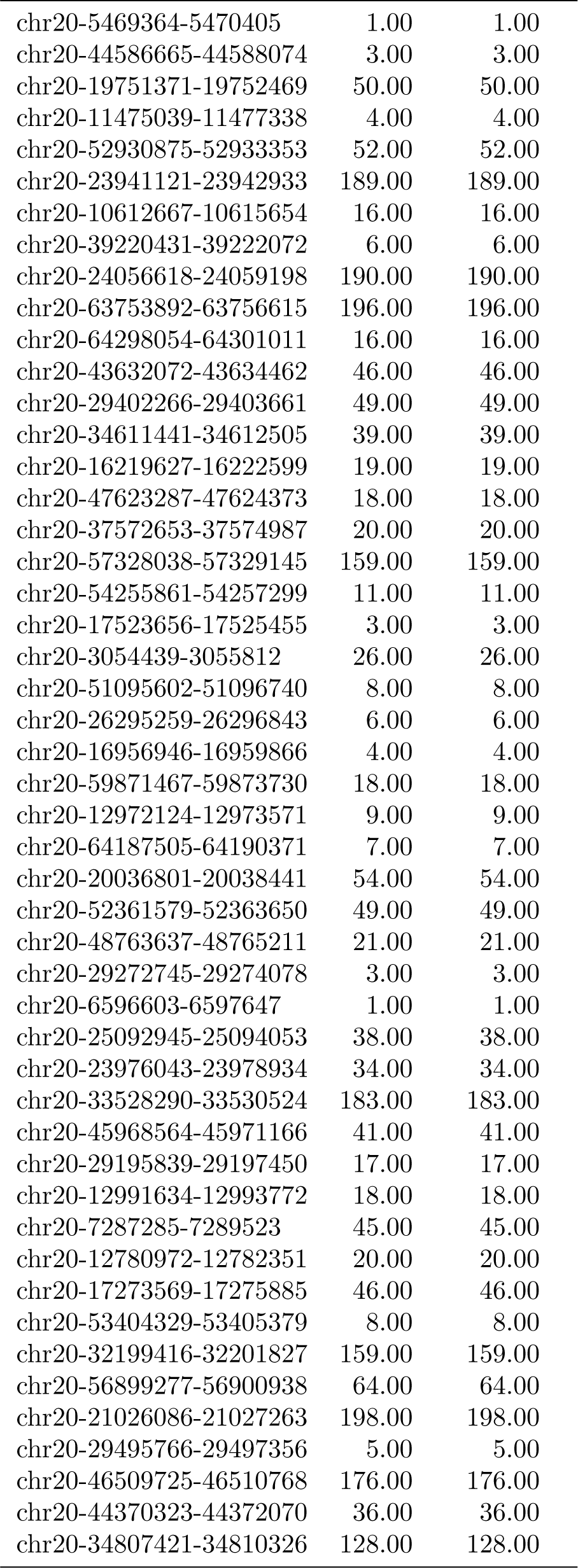
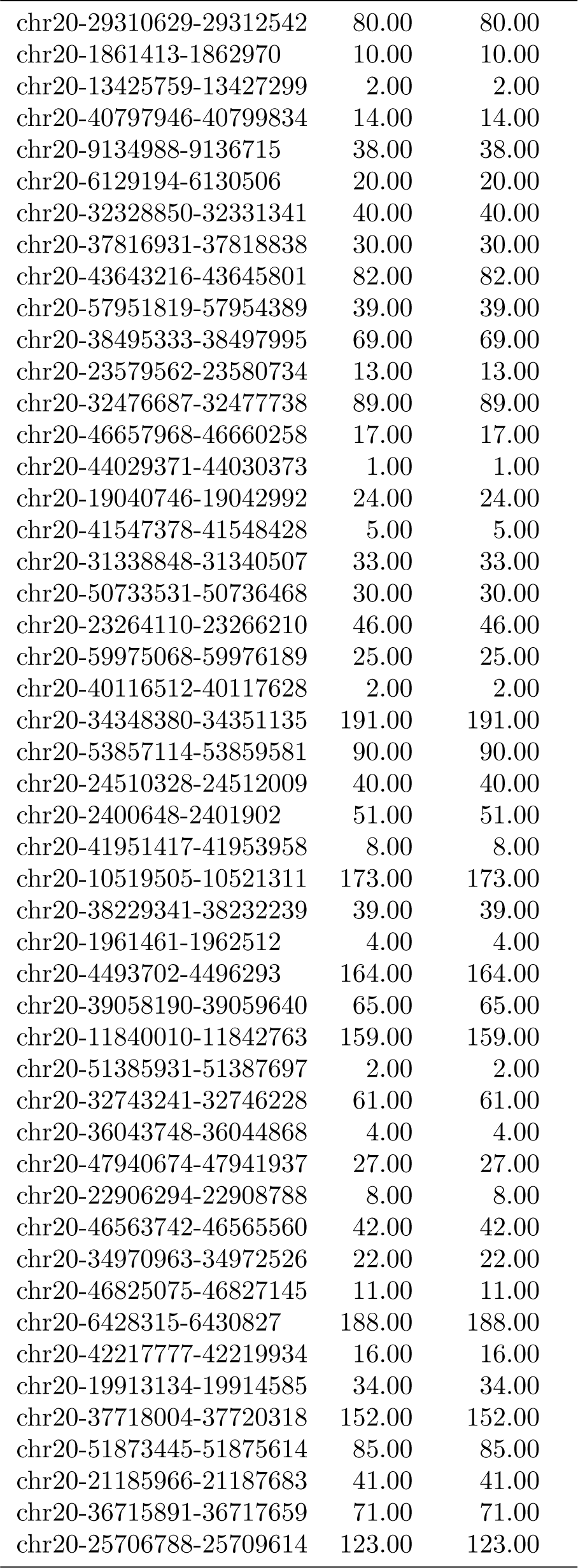
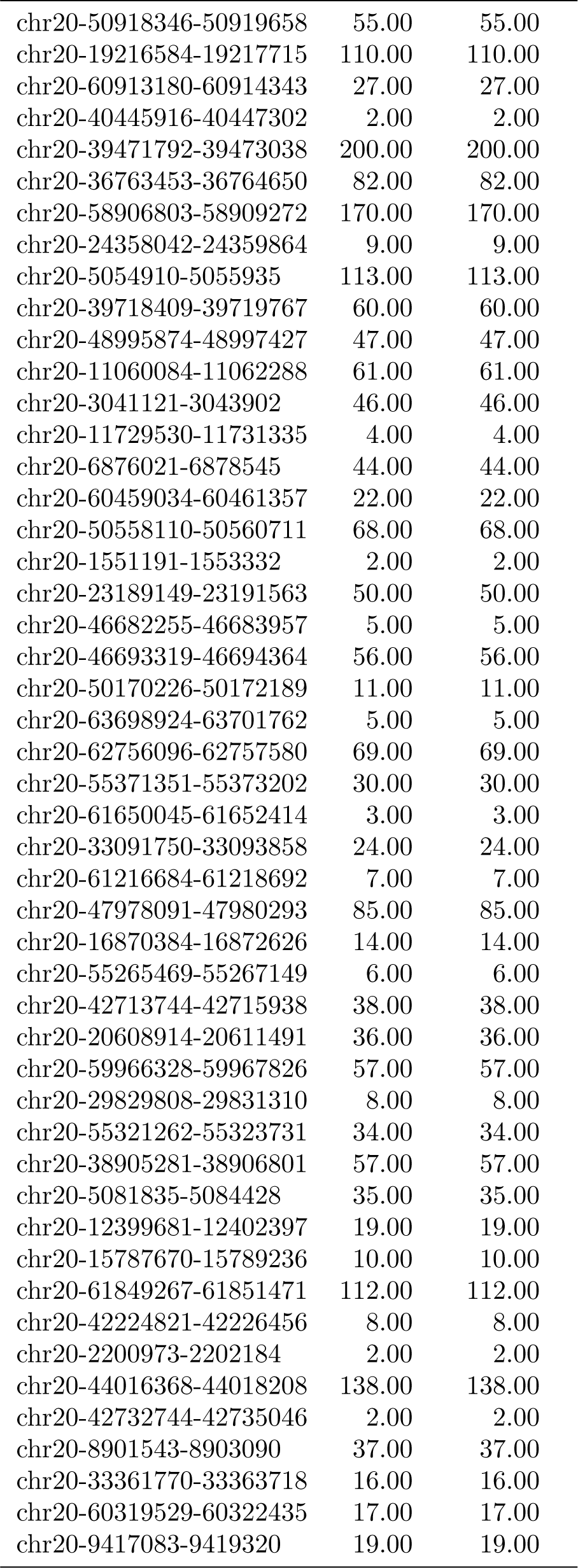
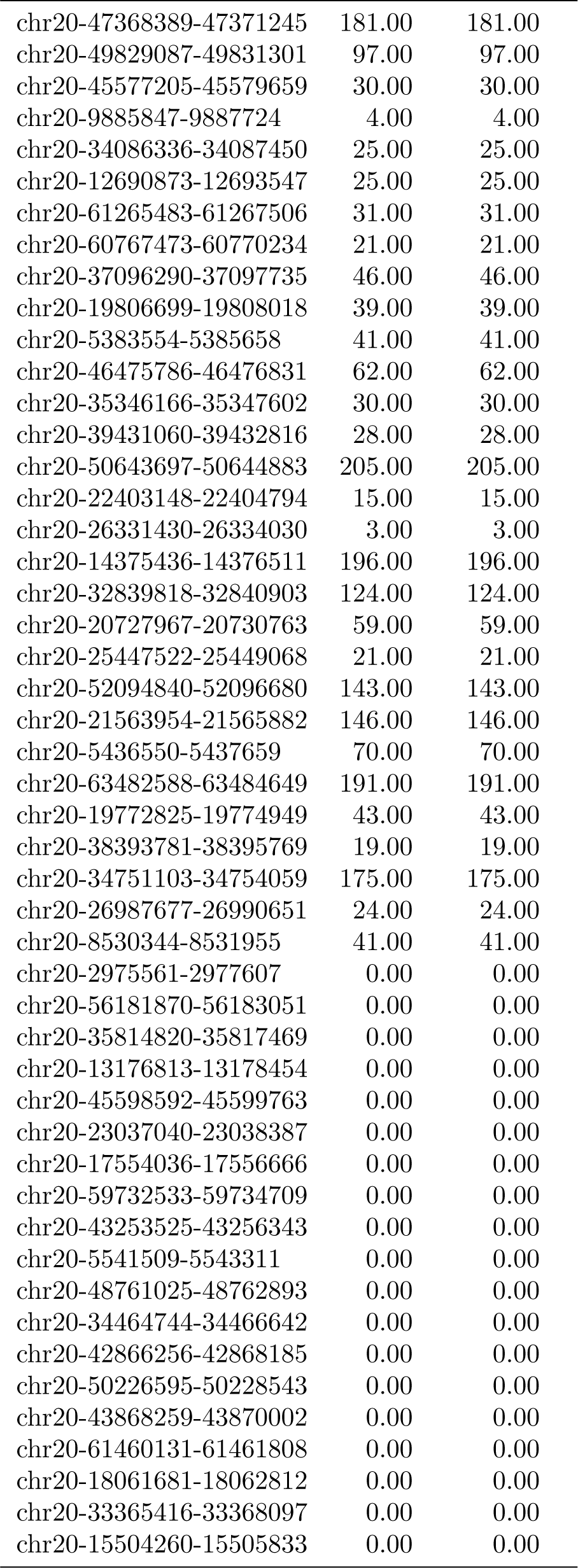
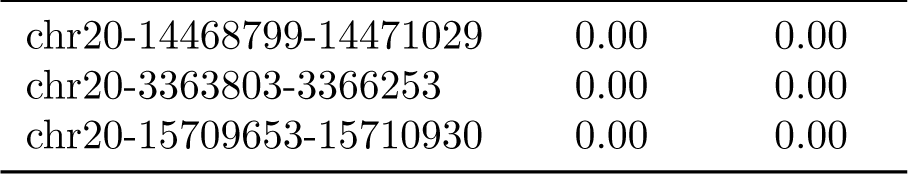
HSP comparison between PxBLAT and BLAT. This table, including 600.00 samples in total, presents a comparison of the HSPs generated by BLAT and PxBLAT for each sample. The column Sample lists the name of the sample, and the columns BLAT and PxBLAT, respectively denote the number of HSPs generated by BLAT and PxBLAT.

| Sample | BLAT | PxBLAT |
| --- | --- | --- |
| chr20-11648866-11650925 | 130.00 | 130.00 |
| chr20-29850079-29852595 | 1.00 | 1.00 |
| chr20-7493878-7496145 | 26.00 | 26.00 |
| chr20-30152074-30153892 | 3.00 | 3.00 |
| chr20-1140095-1141788 | 22.00 | 22.00 |
| chr20-17520690-17523557 | 14.00 | 14.00 |
| chr20-41374266-41376291 | 12.00 | 12.00 |
| chr20-63240773-63243242 | 64.00 | 64.00 |
| chr20-59286740-59288091 | 13.00 | 13.00 |
| chr20-8061734-8063508 | 10.00 | 10.00 |
| chr20-34429296-34431248 | 41.00 | 41.00 |
| chr20-27977914-27979241 | 22.00 | 22.00 |
| chr20-11695455-11697457 | 20.00 | 20.00 |
| chr20-41792348-41794797 | 131.00 | 131.00 |
| chr20-25858004-25860351 | 37.00 | 37.00 |
| chr20-31404034-31407027 | 25.00 | 25.00 |
| chr20-15852904-15855300 | 65.00 | 65.00 |
| chr20-29917054-29919227 | 18.00 | 18.00 |
| chr20-45175162-45177033 | 2.00 | 2.00 |
| chr20-43937342-43939507 | 58.00 | 58.00 |
| chr20-17073859-17076499 | 48.00 | 48.00 |
| chr20-19823346-19824950 | 32.00 | 32.00 |
| chr20-28295597-28298301 | 23.00 | 23.00 |
| chr20-15826504-15829187 | 26.00 | 26.00 |
| chr20-54683378-54684991 | 47.00 | 47.00 |
| chr20-55475241-55476377 | 18.00 | 18.00 |
| chr20-57949934-57952283 | 74.00 | 74.00 |
| chr20-14442978-14444604 | 35.00 | 35.00 |
| chr20-51293808-51295630 | 38.00 | 38.00 |
| chr20-49194155-49196823 | 165.00 | 165.00 |
| chr20-34696351-34697788 | 43.00 | 43.00 |
| chr20-16044321-16045712 | 48.00 | 48.00 |
| chr20-45309757-45312553 | 39.00 | 39.00 |
| chr20-21063232-21065145 | 26.00 | 26.00 |
| chr20-434138-436331 | 174.00 | 174.00 |
| chr20-62110486-62112604 | 167.00 | 167.00 |
| chr20-15866914-15868171 | 15.00 | 15.00 |
| chr20-62343940-62345043 | 69.00 | 69.00 |
| chr20-37168356-37170160 | 34.00 | 34.00 |
| chr20-15699683-15701260 | 15.00 | 15.00 |
| chr20-44826183-44827333 | 152.00 | 152.00 |
| chr20-41538435-41540074 | 43.00 | 43.00 |
| chr20-56032843-56034378 | 18.00 | 18.00 |
| chr20-36813618-36816516 | 21.00 | 21.00 |
| chr20-18637848-18640751 | 68.00 | 68.00 |
| chr20-56294086-56296598 | 27.00 | 27.00 |
| chr20-19514094-19515343 | 2.00 | 2.00 |
| chr20-33893321-33895564 | 126.00 | 126.00 |
| chr20-22282380-22283429 | 4.00 | 4.00 |
| chr20-39910629-39912740 | 84.00 | 84.00 |
| chr20-40898874-40901542 | 27.00 | 27.00 |
| chr20-7536153-7537696 | 15.00 | 15.00 |
| chr20-4818626-4819858 | 60.00 | 60.00 |
| chr20-64316707-64319236 | 24.00 | 24.00 |
| chr20-2960440-2961833 | 119.00 | 119.00 |
| chr20-2519249-2521676 | 46.00 | 46.00 |
| chr20-11596878-11599706 | 68.00 | 68.00 |
| chr20-30829536-30831739 | 16.00 | 16.00 |
| chr20-45188854-45190558 | 3.00 | 3.00 |
| chr20-47238745-47241145 | 14.00 | 14.00 |
| chr20-10538965-10540232 | 83.00 | 83.00 |
| chr20-11996320-11998343 | 87.00 | 87.00 |
| chr20-23440919-23442976 | 47.00 | 47.00 |
| chr20-22451495-22453568 | 4.00 | 4.00 |
| chr20-39709672-39711911 | 23.00 | 23.00 |
| chr20-29782708-29784088 | 8.00 | 8.00 |
| chr20-7911434-7913178 | 9.00 | 9.00 |
| chr20-1508550-1511129 | 193.00 | 193.00 |
| chr20-41451172-41452837 | 6.00 | 6.00 |
| chr20-4003988-4006545 | 29.00 | 29.00 |
| chr20-10921197-10923149 | 5.00 | 5.00 |
| chr20-36789016-36790500 | 186.00 | 186.00 |
| chr20-9309222-9310479 | 157.00 | 157.00 |
| chr20-52532795-52534564 | 9.00 | 9.00 |
| chr20-42081563-42082983 | 4.00 | 4.00 |
| chr20-12605319-12606971 | 2.00 | 2.00 |
| chr20-29389010-29390598 | 149.00 | 149.00 |
| chr20-8128132-8130266 | 170.00 | 170.00 |
| chr20-11399422-11401237 | 12.00 | 12.00 |
| chr20-33251227-33252414 | 2.00 | 2.00 |
| chr20-3342256-3343645 | 92.00 | 92.00 |
| chr20-38869283-38872088 | 60.00 | 60.00 |
| chr20-7649152-7650444 | 60.00 | 60.00 |
| chr20-13021673-13023769 | 58.00 | 58.00 |
| chr20-3971117-3972667 | 32.00 | 32.00 |
| chr20-26232703-26234929 | 61.00 | 61.00 |
| chr20-23063517-23064745 | 7.00 | 7.00 |
| chr20-39079691-39080763 | 26.00 | 26.00 |
| chr20-23882126-23884119 | 92.00 | 92.00 |
| chr20-3170215-3172728 | 44.00 | 44.00 |
| chr20-49421601-49423499 | 35.00 | 35.00 |
| chr20-55380779-55383730 | 204.00 | 204.00 |
| chr20-45794728-45796900 | 164.00 | 164.00 |
| chr20-15159613-15161221 | 9.00 | 9.00 |
| chr20-62378279-62380174 | 184.00 | 184.00 |
| chr20-6934771-6936584 | 116.00 | 116.00 |
| chr20-59538484-59541329 | 8.00 | 8.00 |
| chr20-43661982-43664714 | 26.00 | 26.00 |
| chr20-36707842-36709247 | 85.00 | 85.00 |
| chr20-33898085-33900435 | 65.00 | 65.00 |
| chr20-18062688-18064265 | 46.00 | 46.00 |
| chr20-29490152-29491491 | 23.00 | 23.00 |
| chr20-30806959-30809757 | 0.00 | 0.00 |
| chr20-11837593-11840377 | 40.00 | 40.00 |
| chr20-28934655-28937276 | 1.00 | 1.00 |
| chr20-7689471-7691553 | 18.00 | 18.00 |
| chr20-1746811-1749803 | 9.00 | 9.00 |
| chr20-1261408-1262588 | 16.00 | 16.00 |
| chr20-5757029-5759548 | 29.00 | 29.00 |
| chr20-5361957-5364278 | 107.00 | 107.00 |
| chr20-37305589-37308001 | 21.00 | 21.00 |
| chr20-53702835-53705300 | 148.00 | 148.00 |
| chr20-7447290-7449191 | 16.00 | 16.00 |
| chr20-36626015-36629010 | 150.00 | 150.00 |
| chr20-48184357-48185678 | 2.00 | 2.00 |
| chr20-4634043-4636057 | 9.00 | 9.00 |
| chr20-21292878-21295830 | 179.00 | 179.00 |
| chr20-52551672-52553086 | 7.00 | 7.00 |
| chr20-55410142-55411547 | 55.00 | 55.00 |
| chr20-15164296-15166007 | 21.00 | 21.00 |
| chr20-60294852-60297789 | 197.00 | 197.00 |
| chr20-47112993-47114241 | 38.00 | 38.00 |
| chr20-33645643-33648541 | 62.00 | 62.00 |
| chr20-57099663-57101555 | 91.00 | 91.00 |
| chr20-32443012-32444528 | 1.00 | 1.00 |
| chr20-54597242-54598325 | 31.00 | 31.00 |
| chr20-28168758-28171348 | 20.00 | 20.00 |
| chr20-14255274-14256294 | 211.00 | 211.00 |
| chr20-22858139-22859797 | 8.00 | 8.00 |
| chr20-42977157-42978222 | 11.00 | 11.00 |
| chr20-3890553-3891667 | 46.00 | 46.00 |
| chr20-10043505-10044872 | 33.00 | 33.00 |
| chr20-47292153-47293840 | 50.00 | 50.00 |
| chr20-30056354-30058719 | 0.00 | 0.00 |
| chr20-23406092-23408775 | 15.00 | 15.00 |
| chr20-44245002-44246392 | 36.00 | 36.00 |
| chr20-30999482-31002350 | 1.00 | 1.00 |
| chr20-11990150-11991239 | 3.00 | 3.00 |
| chr20-3810992-3812466 | 197.00 | 197.00 |
| chr20-26419101-26422061 | 0.00 | 0.00 |
| chr20-22091287-22094021 | 119.00 | 119.00 |
| chr20-47477509-47479589 | 59.00 | 59.00 |
| chr20-13750647-13753070 | 44.00 | 44.00 |
| chr20-61483678-61484972 | 2.00 | 2.00 |
| chr20-57584630-57586794 | 13.00 | 13.00 |
| chr20-17904728-17906925 | 34.00 | 34.00 |
| chr20-28086098-28087474 | 19.00 | 19.00 |
| chr20-21317295-21318548 | 51.00 | 51.00 |
| chr20-42758324-42759488 | 193.00 | 193.00 |
| chr20-23580801-23581810 | 23.00 | 23.00 |
| chr20-58929386-58930631 | 22.00 | 22.00 |
| chr20-6605062-6607136 | 79.00 | 79.00 |
| chr20-32987534-32989963 | 169.00 | 169.00 |
| chr20-22493043-22494805 | 22.00 | 22.00 |
| chr20-54190709-54192035 | 24.00 | 24.00 |
| chr20-48343887-48346044 | 2.00 | 2.00 |
| chr20-10262876-10265171 | 189.00 | 189.00 |
| chr20-35374558-35376267 | 67.00 | 67.00 |
| chr20-39018648-39021540 | 83.00 | 83.00 |
| chr20-37785924-37788264 | 202.00 | 202.00 |
| chr20-64334961-64337760 | 0.00 | 0.00 |
| chr20-12575681-12578297 | 14.00 | 14.00 |
| chr20-23374273-23375283 | 1.00 | 1.00 |
| chr20-14278920-14280350 | 40.00 | 40.00 |
| chr20-42778102-42780755 | 9.00 | 9.00 |
| chr20-52810557-52811573 | 83.00 | 83.00 |
| chr20-52564945-52566606 | 22.00 | 22.00 |
| chr20-10598821-10601808 | 32.00 | 32.00 |
| chr20-1172826-1175431 | 87.00 | 87.00 |
| chr20-3347653-3350627 | 44.00 | 44.00 |
| chr20-64176870-64179763 | 43.00 | 43.00 |
| chr20-42829128-42830258 | 100.00 | 100.00 |
| chr20-40623645-40625957 | 187.00 | 187.00 |
| chr20-27009229-27011878 | 24.00 | 24.00 |
| chr20-14732847-14734481 | 95.00 | 95.00 |
| chr20-12698635-12699679 | 2.00 | 2.00 |
| chr20-13328590-13330614 | 13.00 | 13.00 |
| chr20-22912395-22915205 | 12.00 | 12.00 |
| chr20-35339375-35342113 | 48.00 | 48.00 |
| chr20-19431509-19433187 | 28.00 | 28.00 |
| chr20-28727730-28729750 | 16.00 | 16.00 |
| chr20-39515806-39517118 | 2.00 | 2.00 |
| chr20-21857448-21859310 | 209.00 | 209.00 |
| chr20-44751640-44753567 | 19.00 | 19.00 |
| chr20-47371415-47374127 | 45.00 | 45.00 |
| chr20-24140371-24142959 | 6.00 | 6.00 |
| chr20-54083025-54084528 | 41.00 | 41.00 |
| chr20-51775800-51777458 | 46.00 | 46.00 |
| chr20-59400225-59402632 | 30.00 | 30.00 |
| chr20-57144627-57147621 | 47.00 | 47.00 |
| chr20-17690928-17692128 | 6.00 | 6.00 |
| chr20-31238605-31240139 | 4.00 | 4.00 |
| chr20-29259215-29262074 | 4.00 | 4.00 |
| chr20-39132690-39134466 | 34.00 | 34.00 |
| chr20-44674003-44676622 | 31.00 | 31.00 |
| chr20-34435089-34436815 | 13.00 | 13.00 |
| chr20-11085268-11087065 | 27.00 | 27.00 |
| chr20-42010931-42012642 | 10.00 | 10.00 |
| chr20-7483012-7484115 | 1.00 | 1.00 |
| chr20-39873233-39875635 | 207.00 | 207.00 |
| chr20-32029134-32032031 | 93.00 | 93.00 |
| chr20-38052616-38054138 | 32.00 | 32.00 |
| chr20-2602976-2605324 | 19.00 | 19.00 |
| chr20-42169915-42172757 | 15.00 | 15.00 |
| chr20-62125729-62128471 | 19.00 | 19.00 |
| chr20-2162753-2164165 | 5.00 | 5.00 |
| chr20-34630972-34632436 | 113.00 | 113.00 |
| chr20-6611620-6614440 | 152.00 | 152.00 |
| chr20-19300360-19302642 | 6.00 | 6.00 |
| chr20-25189802-25191365 | 154.00 | 154.00 |
| chr20-15951964-15953450 | 26.00 | 26.00 |
| chr20-47285001-47287102 | 52.00 | 52.00 |
| chr20-22548369-22549974 | 9.00 | 9.00 |
| chr20-24725480-24727693 | 52.00 | 52.00 |
| chr20-55633941-55636718 | 5.00 | 5.00 |
| chr20-30324027-30326285 | 30.00 | 30.00 |
| chr20-34789703-34791591 | 27.00 | 27.00 |
| chr20-22886111-22887126 | 5.00 | 5.00 |
| chr20-52373722-52374925 | 52.00 | 52.00 |
| chr20-62594615-62596270 | 61.00 | 61.00 |
| chr20-23567798-23569994 | 5.00 | 5.00 |
| chr20-52746782-52748970 | 26.00 | 26.00 |
| chr20-63919107-63920159 | 20.00 | 20.00 |
| chr20-37603411-37606081 | 21.00 | 21.00 |
| chr20-45549091-45551404 | 167.00 | 167.00 |
| chr20-24122821-24124880 | 2.00 | 2.00 |
| chr20-39384970-39386746 | 47.00 | 47.00 |
| chr20-56953750-56954962 | 11.00 | 11.00 |
| chr20-27265832-27268665 | 27.00 | 27.00 |
| chr20-989008-991475 | 96.00 | 96.00 |
| chr20-13233506-13234845 | 9.00 | 9.00 |
| chr20-30767954-30769153 | 0.00 | 0.00 |
| chr20-54079315-54081921 | 56.00 | 56.00 |
| chr20-22276591-22279281 | 202.00 | 202.00 |
| chr20-32344273-32347087 | 26.00 | 26.00 |
| chr20-14761627-14763228 | 15.00 | 15.00 |
| chr20-24471803-24474607 | 7.00 | 7.00 |
| chr20-22305788-22308754 | 43.00 | 43.00 |
| chr20-63927315-63929626 | 54.00 | 54.00 |
| chr20-33534721-33537439 | 158.00 | 158.00 |
| chr20-53020990-53022920 | 33.00 | 33.00 |
| chr20-44701412-44703325 | 34.00 | 34.00 |
| chr20-63523339-63524524 | 190.00 | 190.00 |
| chr20-39750571-39752199 | 50.00 | 50.00 |
| chr20-47879542-47880820 | 162.00 | 162.00 |
| chr20-12949055-12950079 | 1.00 | 1.00 |
| chr20-37737796-37740734 | 26.00 | 26.00 |
| chr20-17600492-17603407 | 68.00 | 68.00 |
| chr20-40848143-40849212 | 2.00 | 2.00 |
| chr20-33135173-33136880 | 131.00 | 131.00 |
| chr20-39118192-39119260 | 4.00 | 4.00 |
| chr20-27690465-27692366 | 23.00 | 23.00 |
| chr20-37858061-37860321 | 49.00 | 49.00 |
| chr20-12405605-12407882 | 6.00 | 6.00 |
| chr20-57386121-57387852 | 70.00 | 70.00 |
| chr20-34902889-34905611 | 39.00 | 39.00 |
| chr20-52070637-52073331 | 35.00 | 35.00 |
| chr20-40779660-40781751 | 47.00 | 47.00 |
| chr20-31098450-31100717 | 3.00 | 3.00 |
| chr20-29969458-29971235 | 18.00 | 18.00 |
| chr20-13789470-13791842 | 152.00 | 152.00 |
| chr20-26814485-26817257 | 22.00 | 22.00 |
| chr20-40740691-40742983 | 29.00 | 29.00 |
| chr20-49955693-49958425 | 183.00 | 183.00 |
| chr20-37813305-37814584 | 9.00 | 9.00 |
| chr20-18255943-18258048 | 64.00 | 64.00 |
| chr20-63109432-63111043 | 34.00 | 34.00 |
| chr20-14211039-14213272 | 9.00 | 9.00 |
| chr20-17100441-17103140 | 9.00 | 9.00 |
| chr20-2866996-2869328 | 38.00 | 38.00 |
| chr20-53703835-53705398 | 158.00 | 158.00 |
| chr20-51431118-51433743 | 56.00 | 56.00 |
| chr20-6192786-6193790 | 196.00 | 196.00 |
| chr20-10748356-10751019 | 35.00 | 35.00 |
| chr20-33974548-33976407 | 44.00 | 44.00 |
| chr20-32270385-32273111 | 20.00 | 20.00 |
| chr20-36902282-36904506 | 28.00 | 28.00 |
| chr20-198832-201190 | 40.00 | 40.00 |
| chr20-7927314-7928574 | 27.00 | 27.00 |
| chr20-17899461-17900502 | 58.00 | 58.00 |
| chr20-11120841-11122366 | 23.00 | 23.00 |
| chr20-59407661-59409193 | 27.00 | 27.00 |
| chr20-35649824-35652625 | 5.00 | 5.00 |
| chr20-30688628-30690580 | 48.00 | 48.00 |
| chr20-49466264-49467309 | 5.00 | 5.00 |
| chr20-63692672-63694078 | 2.00 | 2.00 |
| chr20-12570747-12573421 | 5.00 | 5.00 |
| chr20-8512304-8514491 | 20.00 | 20.00 |
| chr20-24204303-24205314 | 2.00 | 2.00 |
| chr20-11639430-11641175 | 20.00 | 20.00 |
| chr20-26938330-26940788 | 24.00 | 24.00 |
| chr20-17587799-17589501 | 209.00 | 209.00 |
| chr20-26384682-26386149 | 23.00 | 23.00 |
| chr20-46308421-46310414 | 41.00 | 41.00 |
| chr20-30072951-30074554 | 0.00 | 0.00 |
| chr20-44079162-44081096 | 23.00 | 23.00 |
| chr20-21475551-21476686 | 50.00 | 50.00 |
| chr20-13972488-13974627 | 44.00 | 44.00 |
| chr20-45697903-45699479 | 88.00 | 88.00 |
| chr20-24838480-24839925 | 11.00 | 11.00 |
| chr20-7742386-7745111 | 4.00 | 4.00 |
| chr20-56522851-56524815 | 12.00 | 12.00 |
| chr20-22497057-22499773 | 16.00 | 16.00 |
| chr20-51998729-52000293 | 22.00 | 22.00 |
| chr20-38767581-38768689 | 5.00 | 5.00 |
| chr20-19863311-19865822 | 56.00 | 56.00 |
| chr20-9295230-9297764 | 226.00 | 226.00 |
| chr20-15012779-15015584 | 8.00 | 8.00 |
| chr20-5041846-5044587 | 56.00 | 56.00 |
| chr20-580210-582022 | 35.00 | 35.00 |
| chr20-15328579-15331324 | 42.00 | 42.00 |
| chr20-49481967-49483214 | 17.00 | 17.00 |
| chr20-5260817-5262044 | 5.00 | 5.00 |
| chr20-23340929-23343084 | 102.00 | 102.00 |
| chr20-2269716-2270774 | 189.00 | 189.00 |
| chr20-46399947-46401667 | 31.00 | 31.00 |
| chr20-10815100-10817410 | 32.00 | 32.00 |
| chr20-45864818-45866440 | 53.00 | 53.00 |
| chr20-45489949-45492069 | 35.00 | 35.00 |
| chr20-21302718-21304312 | 1.00 | 1.00 |
| chr20-9931834-9933992 | 5.00 | 5.00 |
| chr20-56295194-56296721 | 5.00 | 5.00 |
| chr20-38225781-38227971 | 18.00 | 18.00 |
| chr20-18585659-18587381 | 35.00 | 35.00 |
| chr20-16940912-16942721 | 39.00 | 39.00 |
| chr20-3758200-3759977 | 4.00 | 4.00 |
| chr20-11807056-11808124 | 3.00 | 3.00 |
| chr20-127575-129488 | 93.00 | 93.00 |
| chr20-40755444-40756701 | 5.00 | 5.00 |
| chr20-21011682-21013735 | 2.00 | 2.00 |
| chr20-10119043-10120171 | 59.00 | 59.00 |
| chr20-1313128-1315640 | 18.00 | 18.00 |
| chr20-11712977-11715178 | 142.00 | 142.00 |
| chr20-34597997-34600726 | 28.00 | 28.00 |
| chr20-53048517-53050101 | 29.00 | 29.00 |
| chr20-15161566-15163488 | 8.00 | 8.00 |
| chr20-15137364-15139959 | 203.00 | 203.00 |
| chr20-60342420-60345056 | 94.00 | 94.00 |
| chr20-55169861-55170982 | 156.00 | 156.00 |
| chr20-24314101-24316265 | 27.00 | 27.00 |
| chr20-53183314-53186148 | 16.00 | 16.00 |
| chr20-43887385-43889661 | 75.00 | 75.00 |
| chr20-59470391-59471545 | 1.00 | 1.00 |
| chr20-53159093-53161302 | 35.00 | 35.00 |
| chr20-5895129-5896312 | 26.00 | 26.00 |
| chr20-25292499-25293803 | 3.00 | 3.00 |
| chr20-34161145-34162369 | 41.00 | 41.00 |
| chr20-45868564-45870103 | 74.00 | 74.00 |
| chr20-4887453-4889888 | 3.00 | 3.00 |
| chr20-39559840-39561722 | 9.00 | 9.00 |
| chr20-33221436-33223116 | 56.00 | 56.00 |
| chr20-755464-758088 | 35.00 | 35.00 |
| chr20-1111176-1112831 | 46.00 | 46.00 |
| chr20-2802607-2803838 | 36.00 | 36.00 |
| chr20-38997152-38999498 | 22.00 | 22.00 |
| chr20-5049946-5052675 | 35.00 | 35.00 |
| chr20-48767423-48769196 | 48.00 | 48.00 |
| chr20-2659893-2661452 | 56.00 | 56.00 |
| chr20-49121260-49124086 | 21.00 | 21.00 |
| chr20-54247731-54250009 | 75.00 | 75.00 |
| chr20-8334685-8337480 | 34.00 | 34.00 |
| chr20-26427340-26428991 | 0.00 | 0.00 |
| chr20-1762478-1764560 | 46.00 | 46.00 |
| chr20-53286968-53289954 | 23.00 | 23.00 |
| chr20-23613637-23615158 | 53.00 | 53.00 |
| chr20-7135001-7136753 | 88.00 | 88.00 |
| chr20-59872605-59875175 | 5.00 | 5.00 |
| chr20-33944307-33946354 | 27.00 | 27.00 |
| chr20-29344297-29347205 | 8.00 | 8.00 |
| chr20-28334593-28336360 | 22.00 | 22.00 |
| chr20-49230825-49232042 | 66.00 | 66.00 |
| chr20-48044237-48045631 | 3.00 | 3.00 |
| chr20-26227485-26229228 | 5.00 | 5.00 |
| chr20-26487906-26489572 | 25.00 | 25.00 |
| chr20-11596974-11599728 | 65.00 | 65.00 |
| chr20-45661385-45663135 | 194.00 | 194.00 |
| chr20-23562380-23564047 | 2.00 | 2.00 |
| chr20-5735736-5738602 | 12.00 | 12.00 |
| chr20-31727237-31729024 | 39.00 | 39.00 |
| chr20-36152090-36154966 | 151.00 | 151.00 |
| chr20-1018906-1020553 | 11.00 | 11.00 |
| chr20-9821106-9823172 | 75.00 | 75.00 |
| chr20-52034533-52037144 | 85.00 | 85.00 |
| chr20-15008181-15010280 | 66.00 | 66.00 |
| chr20-41851737-41853531 | 6.00 | 6.00 |
| chr20-29408974-29411107 | 20.00 | 20.00 |
| chr20-10955881-10957008 | 1.00 | 1.00 |
| chr20-8970270-8971572 | 18.00 | 18.00 |
| chr20-2350436-2352589 | 200.00 | 200.00 |
| chr20-35494115-35496325 | 37.00 | 37.00 |
| chr20-16994461-16996880 | 203.00 | 203.00 |
| chr20-1121638-1123536 | 20.00 | 20.00 |
| chr20-10539946-10542788 | 76.00 | 76.00 |
| chr20-31981394-31984035 | 24.00 | 24.00 |
| chr20-31059406-31062015 | 9.00 | 9.00 |
| chr20-23005241-23006490 | 85.00 | 85.00 |
| chr20-27678668-27680740 | 22.00 | 22.00 |
| chr20-23089029-23090440 | 1.00 | 1.00 |
| chr20-18766639-18768846 | 74.00 | 74.00 |
| chr20-35270491-35271927 | 24.00 | 24.00 |
| chr20-32671767-32673927 | 46.00 | 46.00 |
| chr20-5469364-5470405 | 1.00 | 1.00 |
| chr20-44586665-44588074 | 3.00 | 3.00 |
| chr20-19751371-19752469 | 50.00 | 50.00 |
| chr20-11475039-11477338 | 4.00 | 4.00 |
| chr20-52930875-52933353 | 52.00 | 52.00 |
| chr20-23941121-23942933 | 189.00 | 189.00 |
| chr20-10612667-10615654 | 16.00 | 16.00 |
| chr20-39220431-39222072 | 6.00 | 6.00 |
| chr20-24056618-24059198 | 190.00 | 190.00 |
| chr20-63753892-63756615 | 196.00 | 196.00 |
| chr20-64298054-64301011 | 16.00 | 16.00 |
| chr20-43632072-43634462 | 46.00 | 46.00 |
| chr20-29402266-29403661 | 49.00 | 49.00 |
| chr20-34611441-34612505 | 39.00 | 39.00 |
| chr20-16219627-16222599 | 19.00 | 19.00 |
| chr20-47623287-47624373 | 18.00 | 18.00 |
| chr20-37572653-37574987 | 20.00 | 20.00 |
| chr20-57328038-57329145 | 159.00 | 159.00 |
| chr20-54255861-54257299 | 11.00 | 11.00 |
| chr20-17523656-17525455 | 3.00 | 3.00 |
| chr20-3054439-3055812 | 26.00 | 26.00 |
| chr20-51095602-51096740 | 8.00 | 8.00 |
| chr20-26295259-26296843 | 6.00 | 6.00 |
| chr20-16956946-16959866 | 4.00 | 4.00 |
| chr20-59871467-59873730 | 18.00 | 18.00 |
| chr20-12972124-12973571 | 9.00 | 9.00 |
| chr20-64187505-64190371 | 7.00 | 7.00 |
| chr20-20036801-20038441 | 54.00 | 54.00 |
| chr20-52361579-52363650 | 49.00 | 49.00 |
| chr20-48763637-48765211 | 21.00 | 21.00 |
| chr20-29272745-29274078 | 3.00 | 3.00 |
| chr20-6596603-6597647 | 1.00 | 1.00 |
| chr20-25092945-25094053 | 38.00 | 38.00 |
| chr20-23976043-23978934 | 34.00 | 34.00 |
| chr20-33528290-33530524 | 183.00 | 183.00 |
| chr20-45968564-45971166 | 41.00 | 41.00 |
| chr20-29195839-29197450 | 17.00 | 17.00 |
| chr20-12991634-12993772 | 18.00 | 18.00 |
| chr20-7287285-7289523 | 45.00 | 45.00 |
| chr20-12780972-12782351 | 20.00 | 20.00 |
| chr20-17273569-17275885 | 46.00 | 46.00 |
| chr20-53404329-53405379 | 8.00 | 8.00 |
| chr20-32199416-32201827 | 159.00 | 159.00 |
| chr20-56899277-56900938 | 64.00 | 64.00 |
| chr20-21026086-21027263 | 198.00 | 198.00 |
| chr20-29495766-29497356 | 5.00 | 5.00 |
| chr20-46509725-46510768 | 176.00 | 176.00 |
| chr20-44370323-44372070 | 36.00 | 36.00 |
| chr20-34807421-34810326 | 128.00 | 128.00 |
| chr20-29310629-29312542 | 80.00 | 80.00 |
| chr20-1861413-1862970 | 10.00 | 10.00 |
| chr20-13425759-13427299 | 2.00 | 2.00 |
| chr20-40797946-40799834 | 14.00 | 14.00 |
| chr20-9134988-9136715 | 38.00 | 38.00 |
| chr20-6129194-6130506 | 20.00 | 20.00 |
| chr20-32328850-32331341 | 40.00 | 40.00 |
| chr20-37816931-37818838 | 30.00 | 30.00 |
| chr20-43643216-43645801 | 82.00 | 82.00 |
| chr20-57951819-57954389 | 39.00 | 39.00 |
| chr20-38495333-38497995 | 69.00 | 69.00 |
| chr20-23579562-23580734 | 13.00 | 13.00 |
| chr20-32476687-32477738 | 89.00 | 89.00 |
| chr20-46657968-46660258 | 17.00 | 17.00 |
| chr20-44029371-44030373 | 1.00 | 1.00 |
| chr20-19040746-19042992 | 24.00 | 24.00 |
| chr20-41547378-41548428 | 5.00 | 5.00 |
| chr20-31338848-31340507 | 33.00 | 33.00 |
| chr20-50733531-50736468 | 30.00 | 30.00 |
| chr20-23264110-23266210 | 46.00 | 46.00 |
| chr20-59975068-59976189 | 25.00 | 25.00 |
| chr20-40116512-40117628 | 2.00 | 2.00 |
| chr20-34348380-34351135 | 191.00 | 191.00 |
| chr20-53857114-53859581 | 90.00 | 90.00 |
| chr20-24510328-24512009 | 40.00 | 40.00 |
| chr20-2400648-2401902 | 51.00 | 51.00 |
| chr20-41951417-41953958 | 8.00 | 8.00 |
| chr20-10519505-10521311 | 173.00 | 173.00 |
| chr20-38229341-38232239 | 39.00 | 39.00 |
| chr20-1961461-1962512 | 4.00 | 4.00 |
| chr20-4493702-4496293 | 164.00 | 164.00 |
| chr20-39058190-39059640 | 65.00 | 65.00 |
| chr20-11840010-11842763 | 159.00 | 159.00 |
| chr20-51385931-51387697 | 2.00 | 2.00 |
| chr20-32743241-32746228 | 61.00 | 61.00 |
| chr20-36043748-36044868 | 4.00 | 4.00 |
| chr20-47940674-47941937 | 27.00 | 27.00 |
| chr20-22906294-22908788 | 8.00 | 8.00 |
| chr20-46563742-46565560 | 42.00 | 42.00 |
| chr20-34970963-34972526 | 22.00 | 22.00 |
| chr20-46825075-46827145 | 11.00 | 11.00 |
| chr20-6428315-6430827 | 188.00 | 188.00 |
| chr20-42217777-42219934 | 16.00 | 16.00 |
| chr20-19913134-19914585 | 34.00 | 34.00 |
| chr20-37718004-37720318 | 152.00 | 152.00 |
| chr20-51873445-51875614 | 85.00 | 85.00 |
| chr20-21185966-21187683 | 41.00 | 41.00 |
| chr20-36715891-36717659 | 71.00 | 71.00 |
| chr20-25706788-25709614 | 123.00 | 123.00 |
| chr20-50918346-50919658 | 55.00 | 55.00 |
| chr20-19216584-19217715 | 110.00 | 110.00 |
| chr20-60913180-60914343 | 27.00 | 27.00 |
| chr20-40445916-40447302 | 2.00 | 2.00 |
| chr20-39471792-39473038 | 200.00 | 200.00 |
| chr20-36763453-36764650 | 82.00 | 82.00 |
| chr20-58906803-58909272 | 170.00 | 170.00 |
| chr20-24358042-24359864 | 9.00 | 9.00 |
| chr20-5054910-5055935 | 113.00 | 113.00 |
| chr20-39718409-39719767 | 60.00 | 60.00 |
| chr20-48995874-48997427 | 47.00 | 47.00 |
| chr20-11060084-11062288 | 61.00 | 61.00 |
| chr20-3041121-3043902 | 46.00 | 46.00 |
| chr20-11729530-11731335 | 4.00 | 4.00 |
| chr20-6876021-6878545 | 44.00 | 44.00 |
| chr20-60459034-60461357 | 22.00 | 22.00 |
| chr20-50558110-50560711 | 68.00 | 68.00 |
| chr20-1551191-1553332 | 2.00 | 2.00 |
| chr20-23189149-23191563 | 50.00 | 50.00 |
| chr20-46682255-46683957 | 5.00 | 5.00 |
| chr20-46693319-46694364 | 56.00 | 56.00 |
| chr20-50170226-50172189 | 11.00 | 11.00 |
| chr20-63698924-63701762 | 5.00 | 5.00 |
| chr20-62756096-62757580 | 69.00 | 69.00 |
| chr20-55371351-55373202 | 30.00 | 30.00 |
| chr20-61650045-61652414 | 3.00 | 3.00 |
| chr20-33091750-33093858 | 24.00 | 24.00 |
| chr20-61216684-61218692 | 7.00 | 7.00 |
| chr20-47978091-47980293 | 85.00 | 85.00 |
| chr20-16870384-16872626 | 14.00 | 14.00 |
| chr20-55265469-55267149 | 6.00 | 6.00 |
| chr20-42713744-42715938 | 38.00 | 38.00 |
| chr20-20608914-20611491 | 36.00 | 36.00 |
| chr20-59966328-59967826 | 57.00 | 57.00 |
| chr20-29829808-29831310 | 8.00 | 8.00 |
| chr20-55321262-55323731 | 34.00 | 34.00 |
| chr20-38905281-38906801 | 57.00 | 57.00 |
| chr20-5081835-5084428 | 35.00 | 35.00 |
| chr20-12399681-12402397 | 19.00 | 19.00 |
| chr20-15787670-15789236 | 10.00 | 10.00 |
| chr20-61849267-61851471 | 112.00 | 112.00 |
| chr20-42224821-42226456 | 8.00 | 8.00 |
| chr20-2200973-2202184 | 2.00 | 2.00 |
| chr20-44016368-44018208 | 138.00 | 138.00 |
| chr20-42732744-42735046 | 2.00 | 2.00 |
| chr20-8901543-8903090 | 37.00 | 37.00 |
| chr20-33361770-33363718 | 16.00 | 16.00 |
| chr20-60319529-60322435 | 17.00 | 17.00 |
| chr20-9417083-9419320 | 19.00 | 19.00 |
| chr20-47368389-47371245 | 181.00 | 181.00 |
| chr20-49829087-49831301 | 97.00 | 97.00 |
| chr20-45577205-45579659 | 30.00 | 30.00 |
| chr20-9885847-9887724 | 4.00 | 4.00 |
| chr20-34086336-34087450 | 25.00 | 25.00 |
| chr20-12690873-12693547 | 25.00 | 25.00 |
| chr20-61265483-61267506 | 31.00 | 31.00 |
| chr20-60767473-60770234 | 21.00 | 21.00 |
| chr20-37096290-37097735 | 46.00 | 46.00 |
| chr20-19806699-19808018 | 39.00 | 39.00 |
| chr20-5383554-5385658 | 41.00 | 41.00 |
| chr20-46475786-46476831 | 62.00 | 62.00 |
| chr20-35346166-35347602 | 30.00 | 30.00 |
| chr20-39431060-39432816 | 28.00 | 28.00 |
| chr20-50643697-50644883 | 205.00 | 205.00 |
| chr20-22403148-22404794 | 15.00 | 15.00 |
| chr20-26331430-26334030 | 3.00 | 3.00 |
| chr20-14375436-14376511 | 196.00 | 196.00 |
| chr20-32839818-32840903 | 124.00 | 124.00 |
| chr20-20727967-20730763 | 59.00 | 59.00 |
| chr20-25447522-25449068 | 21.00 | 21.00 |
| chr20-52094840-52096680 | 143.00 | 143.00 |
| chr20-21563954-21565882 | 146.00 | 146.00 |
| chr20-5436550-5437659 | 70.00 | 70.00 |
| chr20-63482588-63484649 | 191.00 | 191.00 |
| chr20-19772825-19774949 | 43.00 | 43.00 |
| chr20-38393781-38395769 | 19.00 | 19.00 |
| chr20-34751103-34754059 | 175.00 | 175.00 |
| chr20-26987677-26990651 | 24.00 | 24.00 |
| chr20-8530344-8531955 | 41.00 | 41.00 |
| chr20-2975561-2977607 | 0.00 | 0.00 |
| chr20-56181870-56183051 | 0.00 | 0.00 |
| chr20-35814820-35817469 | 0.00 | 0.00 |
| chr20-13176813-13178454 | 0.00 | 0.00 |
| chr20-45598592-45599763 | 0.00 | 0.00 |
| chr20-23037040-23038387 | 0.00 | 0.00 |
| chr20-17554036-17556666 | 0.00 | 0.00 |
| chr20-59732533-59734709 | 0.00 | 0.00 |
| chr20-43253525-43256343 | 0.00 | 0.00 |
| chr20-5541509-5543311 | 0.00 | 0.00 |
| chr20-48761025-48762893 | 0.00 | 0.00 |
| chr20-34464744-34466642 | 0.00 | 0.00 |
| chr20-42866256-42868185 | 0.00 | 0.00 |
| chr20-50226595-50228543 | 0.00 | 0.00 |
| chr20-43868259-43870002 | 0.00 | 0.00 |
| chr20-61460131-61461808 | 0.00 | 0.00 |
| chr20-18061681-18062812 | 0.00 | 0.00 |
| chr20-33365416-33368097 | 0.00 | 0.00 |
| chr20-15504260-15505833 | 0.00 | 0.00 |
| chr20-14468799-14471029 | 0.00 | 0.00 |
| chr20-3363803-3366253 | 0.00 | 0.00 |
| chr20-15709653-15710930 | 0.00 | 0.00 |

## Acknowledgments

Special thanks to the team who maintain the UCSC codebase and users from the bioinformatics community whose valuable feedback and suggestions were pivotal in refining PxBLAT’s design and functionality.

